# Charged agar surfaces affect *E. coli* biofilm properties by balancing curli amyloid quantity and quality

**DOI:** 10.64898/2026.04.27.721109

**Authors:** Macarena Siri, Mónica Vázquez-Dávila, Cécile M. Bidan

## Abstract

Biofilm extracellular matrix (ECM) varies with environmental conditions and substrate properties. Understanding the surface-biofilm relationship helps to perfect antibacterial strategies and to design new engineered living materials (ELMs). In this work, we studied how cationic and anionic polyelectrolyte coatings affect macroscopic features of *Escherichia coli* curli-producing biofilms, as well as the properties of their curli amyloid fibers. Cationic coatings limited biofilm spreading, increased their surface density and water absorption, which correlated with a higher yield of curli amyloid fibers with looser structure. In contrast, anionic surfaces allowed for standard biofilm spreading, with a lower fiber yield but a more compact and chemically stable fiber structure. Higher biofilm rigidity and adhesion were measured on both types of charged surfaces. Thus, we propose that the differences in biofilm macroscopic properties result from a trade-off between curli quantity and quality in the ECM, namely fiber density and molecular packing, as well as their interaction with water. Our findings provide insights on how the biophysical properties of the ECM can be controlled by tuning the substrate physico-chemical characteristics with charged coatings. This work opens up new avenues for developing antimicrobial strategies, as well as tailoring the properties of amyloid-based ELMs.

**TOC figure:** 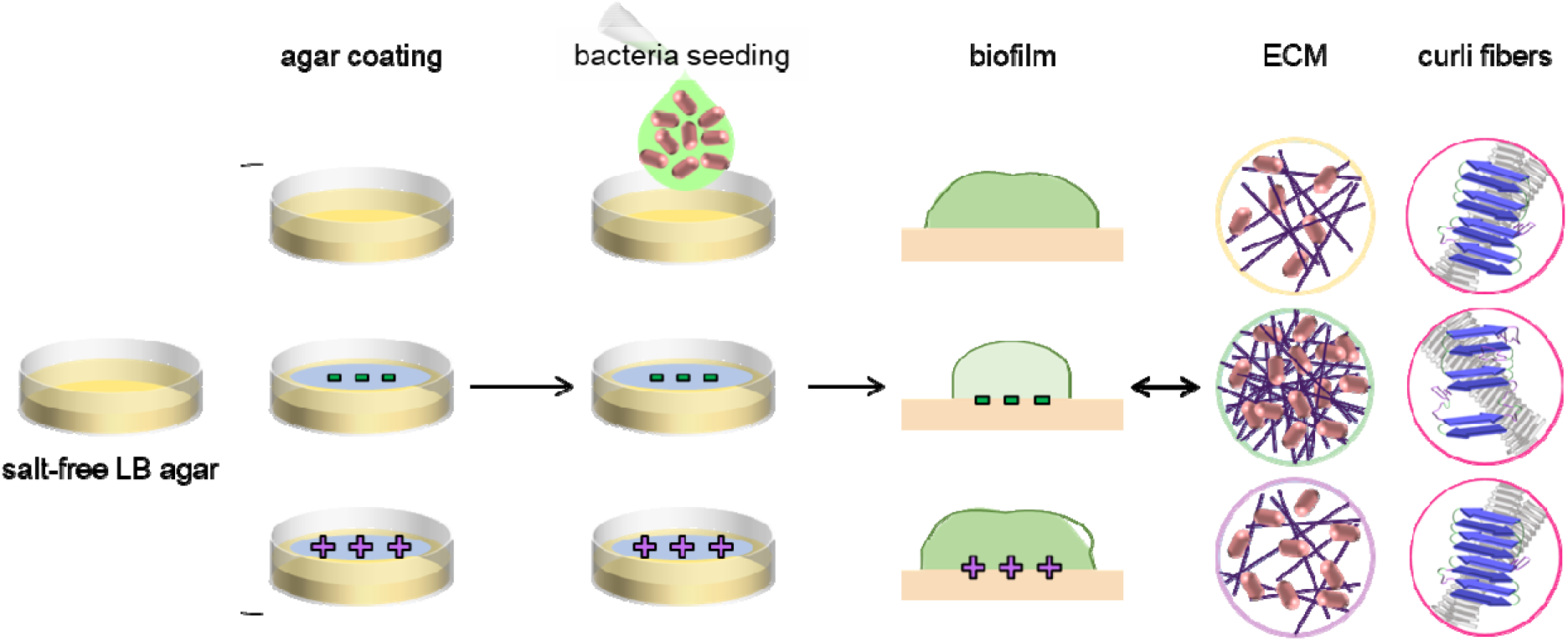

## Introduction

Bacteria attach to surfaces, gather in colonies and get embedded in a self-produced polymeric matrix to form biofilms.^1,2^ This prevalent bacterial form of life is sessile, cooperative and enhances resistance to mechanical damages and antibiotic agents. These advantages are provided by the extracellular matrix (ECM), which consists of a network of biopolymers, mainly made of polysaccharides, proteins, and nucleic acids.^3,4^ The composition and quantity of the ECM vary with the environmental conditions (e.g. levels of oxygen and/or nitrogen, temperature, pH, availability of nutrients and/or water) and the properties of the underlying surface.^5^ Understanding these relationships is of interest to develop strategies aiming at weakening biofilms in the quest of antibacterial materials,^6^ or at tuning biofilm functional properties for their use in engineered living materials (ELMs).^7^

Over the years, research has focused on how ECM properties contribute to biofilm response to environmental cues. For example, the secretion of exopolysaccharides in biofilms drives surface motility by generating osmotic pressure gradients in the extracellular space.^8^ Alongside, it was proposed that for biofilms containing protein amyloid fibers, the amyloid assembly reaction is likely to represent a means for controlling biofilm formation.^9^ In *P. aeruginosa*, Fap amyloids contribute to biofilm rigidity.^10^ In *B. subtilis* biofilms, TasA protein forming amyloid fibers determine biofilm morphology and robustness.^11^ Further studies have focused on how environmental cues affect biofilm development. In *Vibrio cholerae* biofilms, matrix production enables biofilm-dwelling bacterial cells to establish an osmotic pressure difference between the biofilm and the external environment as a way of enhancing nutrient uptake.^1^ In this study, there was a synergistic role of osmotic pressure and crosslinking of the matrix to control biofilm growth and to maintain its protective functions.^1^ Nutrient availability and salinity also contribute to the morphology, and growth kinetics of *V. fischeri* biofilms.^12^ Recently, few studies have deepened the understanding of how the interplay across scales within the biofilm influence their adaptability and survival. In *E. coli* biofilms, agar water content influences not only the spreading and morphology of the biofilms,^3,13^ but also the structure and function of curli amyloid fibers, which in turn contribute to biofilm rigidity.^3^ As the structure of curli fibers loosen upon water availability, their chemical stability decreases together with biofilm rigidity. Likewise, nutrient availability was shown to impact bacterial protein expression, the structure and function of curli amyloid fibers, and biofilm mechanical properties.^4^

In addition to environmental cues, surface properties of biofilm substrate (e.g., surface energy) and electrostatic interactions between substrate and bacteria also influence biofilm formation.^14,15^ When occurring at an interface, biofilm development is determined by the initial adhesion of single bacteria to the surface, and the further biofilm growth and spreading due to cell proliferation and biopolymer production.^15^ Physicochemical properties such as charge, hydrophobicity and chemistry of the substrate have an impact on the initial attachment of bacteria.^16^ The biological production of ECM results in an osmotic flux of water from the agar into the biofilm that subsequently swells and spreads out.^14^ Cationic coatings for instance, limit biofilm spreading, but promote biofilm density in *Escherichia coli* depending on the degree of polyelectrolyte protonation.^15^ The architecture of *P. aeruginosa* ECM varies with the material properties of the substrate. The mesh size of the hydrogel contributes to the bacteria distribution in the early stages of biofilm development, and high-density substrates form shallow biofilms.^16^ Polyelectrolyte coatings change the physical and mechanical characteristics of agar substrates and are thus used as antimicrobial agents to prevent the growth of unwanted biofilms.^17^ Despite this, the influence of charged surfaces on bacterial ECM and their consequences on the emergent biofilm properties are still not fully understood.

In this work we studied the adaptation of amyloid-producing *E. coli* biofilms (W3110) to polyelectrolyte-modified agar gels at the biofilm level (mechanical properties), and at the molecular level (structure and function of curli amyloid fibers). Our results showed that a balance between quantity and quality of curli amyloid fibers determines biofilms macroscopic characteristics.

## Results

### 1- Surface charge affects ECM production and interaction with water

Following up on Ryzhkov et al.^15^, we studied the effect of the coating of nutritive substrates with two polyelectrolytes - polyethyleneimine (PEI; weak polycation) and poly (acrylic acid) (PAA; weak polyanion) – on *E. coli* W3110 biofilms and their curli amyloid ECM fibers. After 5 days of growth, we observed differences between the morphology of PEI-grown *E. coli* biofilms and the biofilms on non-modified surfaces used as reference (Figure 1a, and Figure S1). While the reference biofilms were translucent and presented concentric wrinkling (observed as brighter areas)^4,18,19^, PEI-grown biofilms were more opaque, devoid of these brighter areas, and presented a non-round shape. PAA-grown biofilms were similar to the reference biofilms, but no brighter areas were observed^15^. As expected, biofilms grown on PEI occupied a third of the area of *E. coli* W3110 biofilms grown on non-modified surfaces (92 ± 1 and 314 ± 7 mm^2^, respectively), while no significant differences were found between the area of the control biofilms, and the biofilms grown on PAA-modified surfaces (c.a. 280 ± 30) (Figure 1b).^15^ We estimated the dry mass of the biofilms produced in each condition. While PEI-grown biofilms presented half the dry weight of the reference, there were no significant differences between the control and PAA-grown biofilms (3 ± 0.1, 6 ± 0.3, and 5 ± 0.5 mg, respectively) (Figure 1a). We define the surface density (δ) as the biofilm dry mass per mm^2^ of the agar surface occupied by the biofilm (Figure 1b). While PAA-grown biofilms had similar surface density values to the reference biofilms (0.02 ± 0.0005 and 0.02 ± 0.0001 mg/mm^2^, respectively), PEI-grown biofilms almost doubled these values with a surface density of 0.03 ± 0.0007 mg/mm^2^.

**Figure 1.**
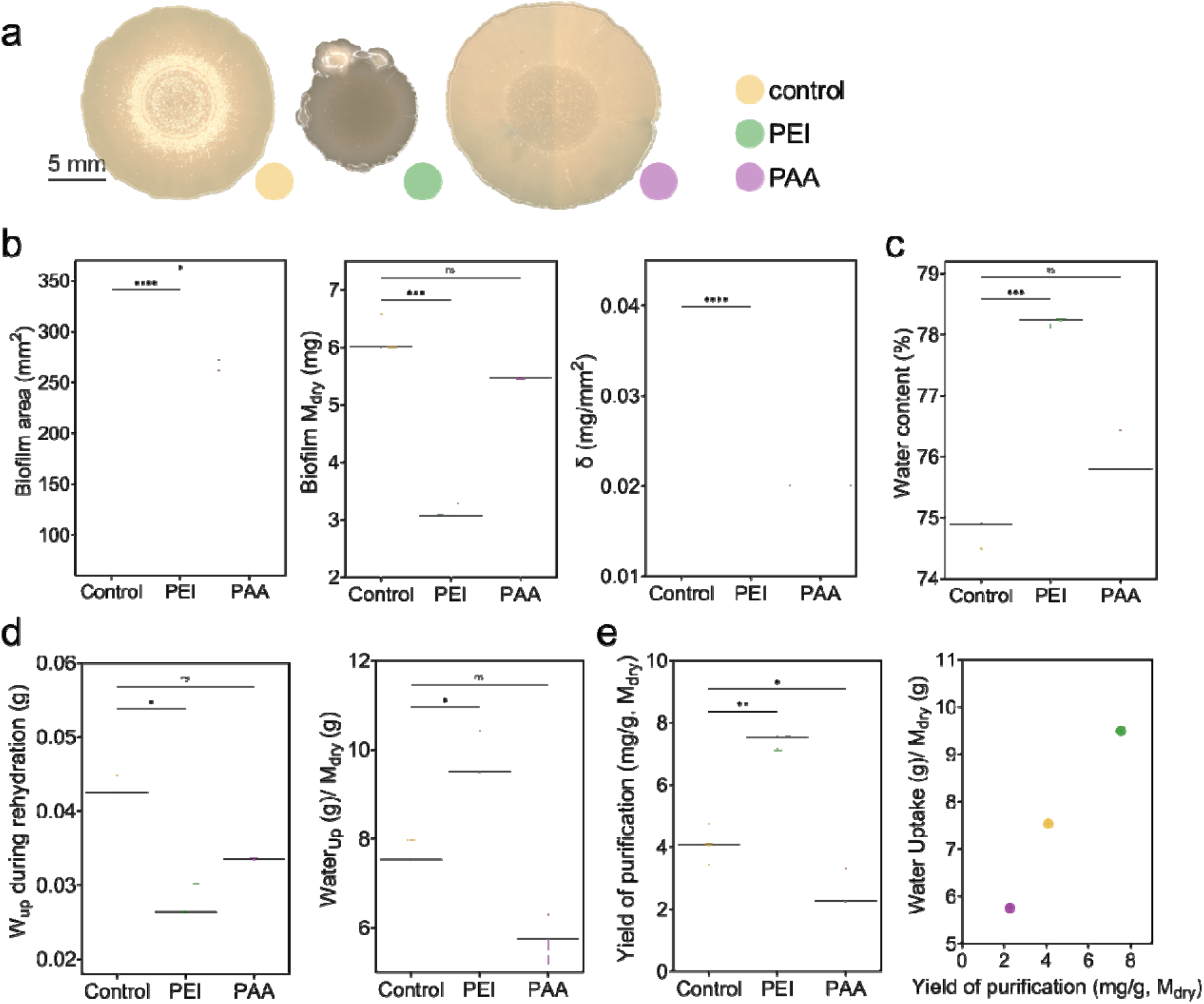
General characteristics of *E. coli* W3110 biofilms grown on salt-free LB-agar substrates with different coatings. (a) Representative phenotype of five-day old biofilms grown on salt-free LB-agar substrates with different coatings. (b) Biofilm area (n= 7-10 independent biofilms), biofilm dry mass (M_dry_) and biofilm density (δ) as a function of composition (n= 7-10 independent biofilms). Median, quartiles and extreme values are represented by the line, the box limits and the whiskers respectively. The statistical analysis was done with Man Whitney U test (p<0.0001, **** | p<0.001, *** | p<0.01, ** | p<0.05, | ns = non-significant). (c) Water content of biofilms. (d) Water uptake (W_up_) per biofilm after overnight rehydration in excess of water and water uptake per M_dry_. (e) Purification yield of the curli fibers extraction process in milligram of CsgA per gram of biofilm dry mass. The CsgA content in the purified curli fibers was estimated by absorbance after denaturation using 8 M urea. Correlation of the W_up_ per M_dry_ of biofilm and the yield of purification of the fibers per M_dry_. Unless stated otherwise, all data presented here come from N=4 independent biofilm cultures for each condition tested, and the statistical analysis was done with One-way ANOVA (p<0.001, *** | p<0.01, ** | p<0.05, * | ns = non-significant). The control condition was used as reference.

PEI-grown biofilms had the highest water content (78 ± 0.14 %) (Figure 1c). While the rehydration capacity of these biofilms was the lowest (c.a. 0.03 ± 0.002 g), when relativized to the biofilm dry mass the PEI-grown biofilms showed the highest water absorption capacity (9 ± 1 g/g) (Figure 1d). This property may be related to the high curli amyloid fiber yield obtained after extraction from PEI-grown biofilms (7 ± 0.4 mg/g) (Figure 1e). While these biofilms had the highest fiber yield per biofilm dry mass, PAA-grown biofilms showed the lowest fiber yield per biofilm dry mass (2 ± 1 mg/g), reducing by half the fiber yield compared to biofilms grown on non-modified surfaces (c.a. 4 ± 0.6 mg/g). Results so far suggest that surface properties of the agar such as the charge affect bacterial extracellular matrix production, biofilm water absorption, as well as biofilm spreading properties.

### 2- Surface coating yields stiffer biofilms, less plastic and more adhesive

We studied the mechanical properties of the biofilms with microindentation including biofilm stiffness (reduced elastic modulus, E_r_), apparent plasticity index (Ψ’), and adhesion force between the indentation tip and the biofilm surface (F_ad_) (Figure 2, Figure S2).^4,19,20^ Biofilm reduced elastic modulus was calculated by fitting the first 10 μm of indentation to a Hertzian contact model equation (see Experimental section for details) (Figure 2a). *E. coli* biofilms grown on a non-modified surface had the lowest reduced modulus median values of 14 kPa, while the PAA-grown, and PEI-grown biofilms presented a stiffness of 54 and 134 kPa, respectively. Interestingly, biofilm rigidity measured in the three growth conditions appeared to increase with their water content (Figure 2b), which seems counterintuitive as diluting the polymers of a hydrogel with water is expected to decrease its mechanical properties.^21^

**Figure 2.**
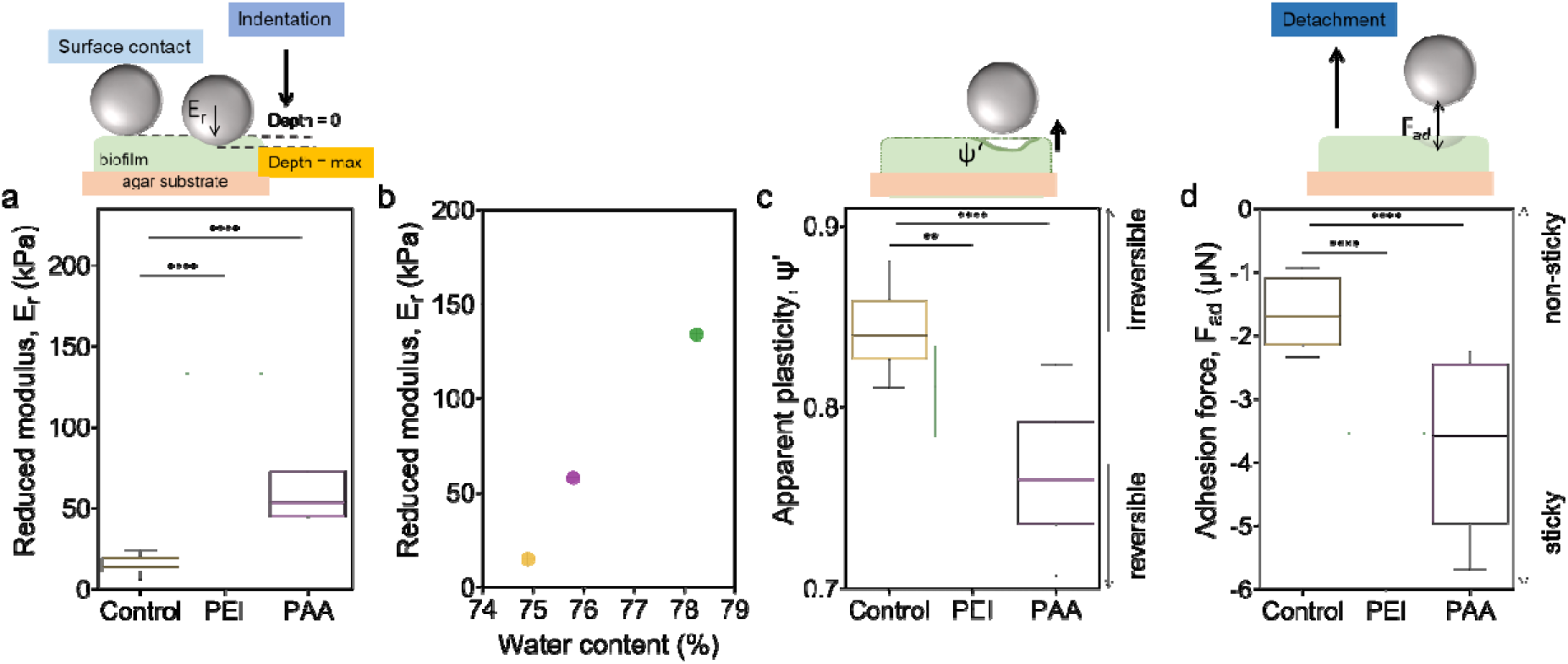
Mechanical characterization of the different strains of *E. coli* biofilms with microindentation. (a) Averaged reduced elastic modulus performed on biofilm surfaces (10 individual measurements per biofilm, with 3 independent biofilms per strain). (b) Correlation between biofilm reduced elastic modulus, (E_r_) and biofilm water content (%). (c) Apparent plasticity index (ψ’) describes the capacity of the biofilm to deform irreversibly when a load is applied. (d) Adhesion force, F_ad_ measured from the minimum load recorded during tip retraction divided by the contact area at maximum indentation. Except from (a), data obtained correspond to indentation curves with a maximum depth of 20 μm. N = 3 different biofilms. The statistical analysis was done with Man Whitney U test (p<0.0001, **** | p<0.001, *** | p<0.01, ** | p<0.05, * | ns = non-significant). A minimalistic sketch was included on top of each plot to highlight the phenomenon focused on in each analysis. A minimalistic sketch was included on top of each plot to illustrate the phenomenon probed in each analysis.

We define the apparent plasticity index (Ψ’) to characterize the irreversibility of the deformation induced by the indentation (Figure 2c).^3^ Higher Ψ’ values indicate irreversible changes in the biofilm upon loading (plastic behavior). Even though all biofilms presented a Ψ’ median value around 0.8, biofilms grown on non-modified surfaces and PEI-grown biofilms presented the highest Ψ’ with 0.84 and 0.81, respectively. PAA-grown biofilms presented a plastic Ψ’ index of 0.76.

We determined the adhesion force (F_ad_) at which the indentation tip starts to detach from the biofilm during its retraction.^4^ The lower the value of the F_ad_, the higher the force needed for tip detachment. Biofilms grown on non-modified surfaces had the highest (less negative) F_ad_ median values (-1.7 μN), while the PEI-grown and PAA-grown biofilms more negative (lowest) F_ad_ median values at -3.5 μN. This suggests that adhesion to the surface of biofilms grown on charged surfaces was higher than to biofilms grown on non-modified surfaces (Figure 2d).

Results suggest that polyelectrolyte coatings change the mechanical properties of *E. coli*. When grown on charged surfaces, the biofilms become stiffer, more plastic and stickier than when grown on non-modified surfaces.

### 3- Polycation surface coating makes E. coli curli fibers less structured

Charged-coated surface proved to change the macroscopic characteristics of the biofilms; not only the biofilm morphology (Figure 1a), but also their interaction with water (Figure d-e), and mechanical properties (Figure 2). To have an insight of how these coatings influenced the ECM, we purified the curli amyloid fibers from the biofilms. After extraction from the biofilms grown in different conditions, the curli amyloid fibers were observed with Transmission Electronic Microscopy (TEM) to confirm the existence of fibers (Figure 3a). The images also showed rough differences in their morphology. Curli extracted from PEI-grown biofilms appeared more tube-like than the curli fibers extracted from biofilms grown on non-modified agar surfaces. PAA-grown biofilms produced similar fibers to the curli fibers acquired on non-modified substrates.

**Figure 3.**
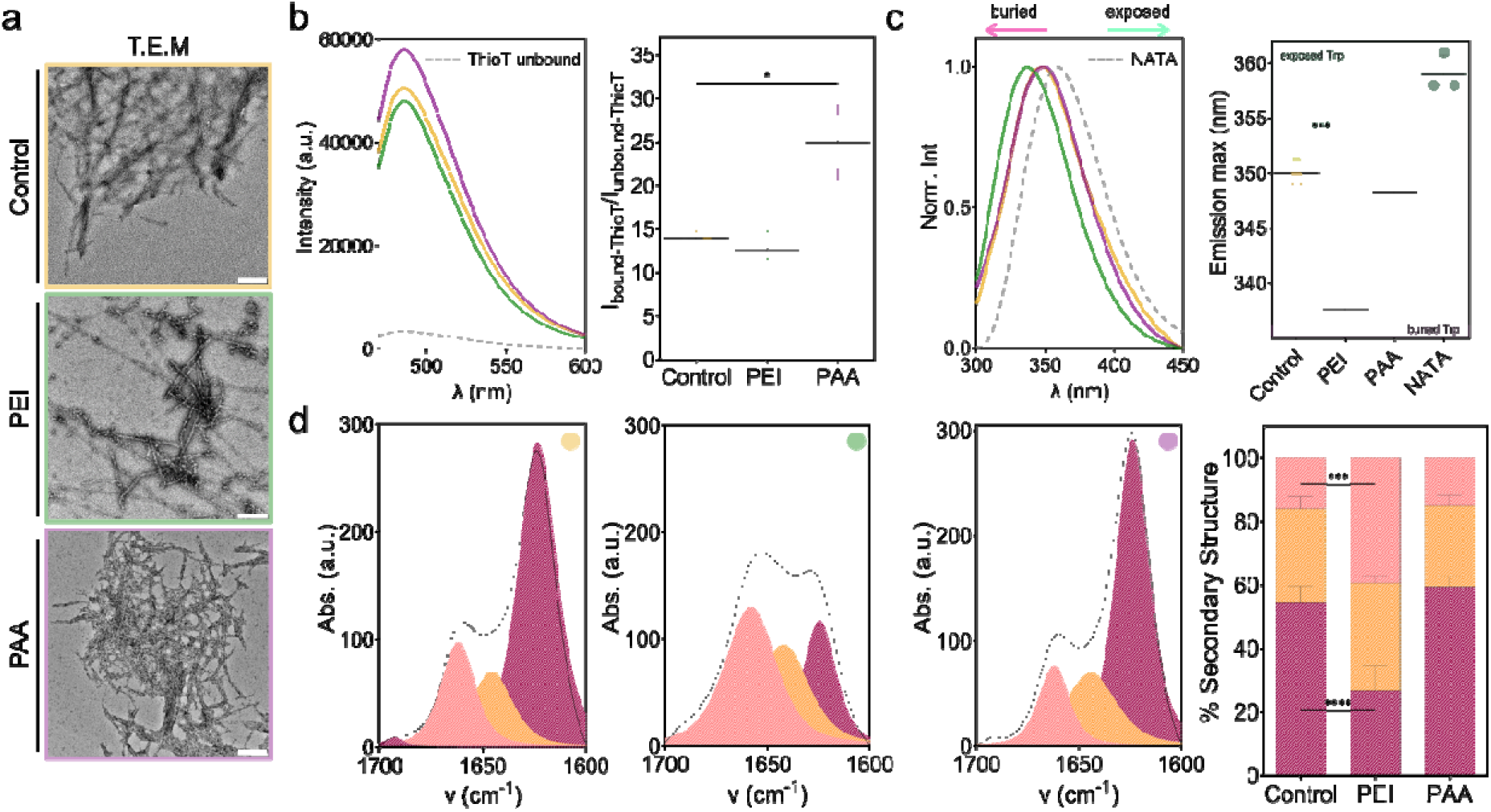
Microscopy and spectroscopic study of the structure of the purified curli amyloid fibers. (a) Electron microscopy images of the purified fibers from the different biofilms (scale bar= 500 nm). (b) ThioT fluorescence emission spectra of the fibers after purification (left panel), and the ratio between bound and unbound ThioT (right panel). (c) Representative intrinsic fluorescence spectra of the fibers Trp emission. The spectrum of soluble Trp (NATA) in buffer is represented as reference for the maximum exposure possible of the Trp to the surface (dotted line) (λexc = 280 nm). Position of the emission peak of the intrinsic fluorescence of the fibers through Trp emission (right panel). The information was extracted from spectra like the ones represented in the left panel. The spectrum of soluble Trp (NATA) in buffer (dotted line) is represented as reference for the maximum exposure possible of the Trp to the surface (λexc = 280 nm). Areas shadowed in pink or green indicate the extreme positions for hydrophobic or hydrophilic Trp, respectively. (d) Representative ATR-FTIR spectra of the Amide I’ region of each fiber purified from the different biofilms (left panels). Distribution of the secondary structure of the curli fibers studied. The data was obtained from the Amide I’ region of each spectra (right panel). The statistical analysis was done with One-way ANOVA, post test used was Tukey’s test to compare each fiber against every fiber (p < 0.0001, **** | p < 0.001, *** | p < 0.01, ** |p < 0.05, * | ns = non-significant). For all experiments, N=3 independent purification batches of fibers grown in each condition.

Thioflavin T (ThioT) is a probe used to identify amyloid fibers, having affinity for β-sheet enriched structures.^5,6^ All fibers had ThioT emission indicating the presence of β-sheet rich structures (Figure 3b). While the emission of ThioT bound to fibers extracted from PEI-grown biofilms did not show significant differences with the emission of ThioT bound to control fibers, fibers from PAA-grown biofilms presented a significantly higher ThioT emission. Differences in the ratio between the ThioT bound to fibers and ThioT unbound signal suggests a difference regarding β-sheet, either in the amount and/or packing of these structures in the fibers.^7,8^ Figure 3b suggests that fibers from PAA-grown biofilms have higher β-sheet content, or that the spacing between these structures is lower than in the other fibers tested.

The CsgA monomer that forms curli fibers contains a single tryptophan (Trp) in its composition. Trp serves as a solvatochromic fluorophore, where the maximum emission wavelength gives information on the hydrophobicity of the Trp environment, thus indicating changes in the protein structure (Figure 3c).^7–9^ We used soluble Trp (NATA) as reference for the maximum exposure position. Curli amyloid fibers from PEI-grown biofilms presented a blueshift in their Trp population emission spectrum, while curli fibers from PAA-grown biofilms did not present any difference of Trp position when compared to fibers from biofilms grown on non-modified surfaces. The peak position in the curli fibers from PEI-grown suggested a Trp population buried deep in the fiber structure (337 ± 1 nm), while the Trp population in fibers from biofilms grown on non-modified agar surfaces and from PAA-grown biofilms were exposed to the solvent (350 ± 1, and 348 ± 3 nm, respectively).

Further structure differences of the fibers were found by ATR-FTIR spectroscopy experiments (Figure 3d, Table1). We focused on the Amide I’ region (1700 – 1600 cm^-1^) as it is the most susceptible to structure changes.^7,8,10,11^ The spectrum of the fibers from biofilms grown on non-modified agar surfaces reported the typical AT - 1620 cm^-1^ the spectrum presented a sharp band assigned to β-sheet structures, and a second band around 1660 cm^-1^ assigned to β-turn structures.^11^ Fibers from PAA-grown biofilms presented a similar spectrum, whereas the spectrum of fibers from PEI-1660 cm^-1^ 1620 cm^-1^. The peak inversion suggested a larger β-turn content than β-sheet content in the PEI-grown fibers. Moreover, the difference between spectra suggests a profound change in the fibers structure with a decrease in β-sheet content by half from 54 % to 27 % and an increase in β-turn content from 16 % to 40 % of the fibers from PEI-grown biofilms compared to fibers from biofilms grown on non-modified agar surfaces (Table 1). These results on the structure of the fibers suggest that the polycation-coated surface influences curli amyloid fibers formation in these biofilms.

**Table 1.** Secondary structure content of the curli fibers. Data was extracted from Figure 3.

| Secondary structure % | Control | PEI | PAA |
| --- | --- | --- | --- |
| Beta-sheet | 54.28 ± 5.33 | 26.66 ± 7.79 | 59.48 ± 3.09 |
| Random/alpha | 29.63 ± 3.84 | 33.89 ± 2,18 | 25.35 ± 3.34 |
| Beta-turns | 16.08 ± 1.63 | 39.45 ± 8.46 | 15.16 ± 0.36 |

**Table 2.**
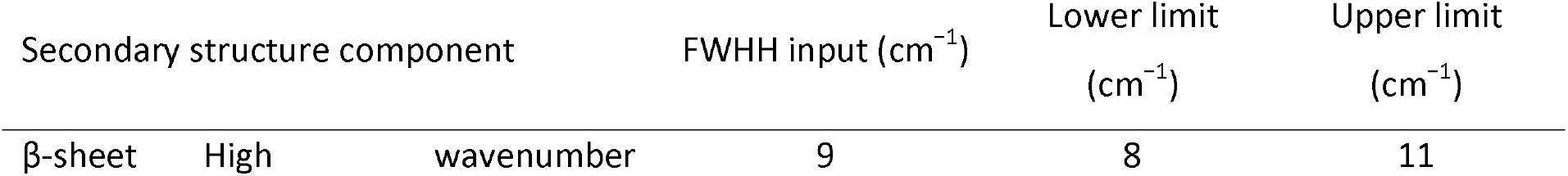
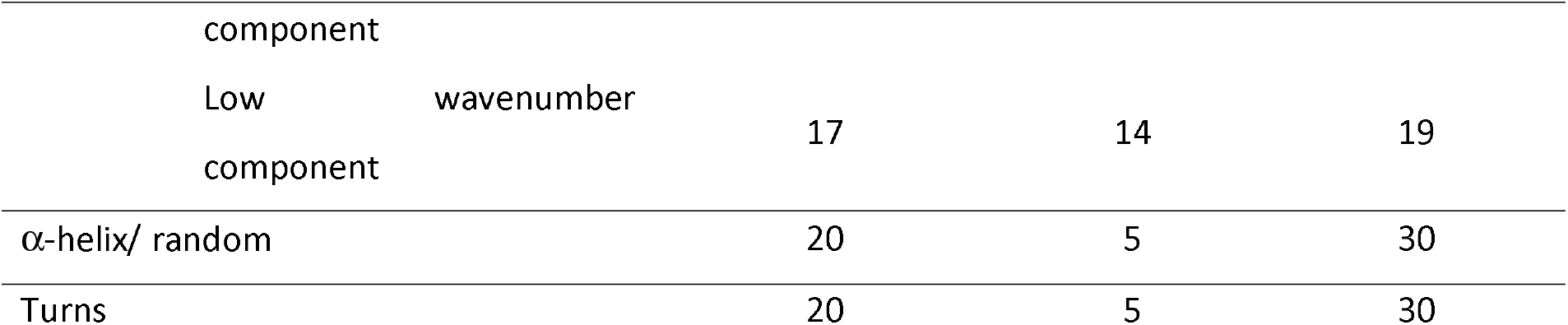
FWHH input values in cm−1 and their physically plausible ranges expected for each type of secondary structure ^17–19^.

### 4- Polycation surface coating makes E. coli curli fibers more chemically unstable

Finally, in order to assess the consequences of the changes in the fiber structure, we studied their hydrophobic character, and chemical stability (Figure 4). Nile Red (NR) is a solvatochromic probe used to characterize the hydrophobicity of molecular structures such as amyloid fibers.^7,8,12^ In contrast to the Trp, NR is used here as an external probe to report differences in the hydrophobicity of the fibers studied. Curli fibers from biofilms grown on non-modified surfaces presented the most hydrophilic character, while the fibers from PEI-grown biofilms presented the most hydrophobic character (Figure 4a). This result follows the trend observed when the Trp intrinsic hydrophobicity was studied (Figure 3c).

**Figure 4.**
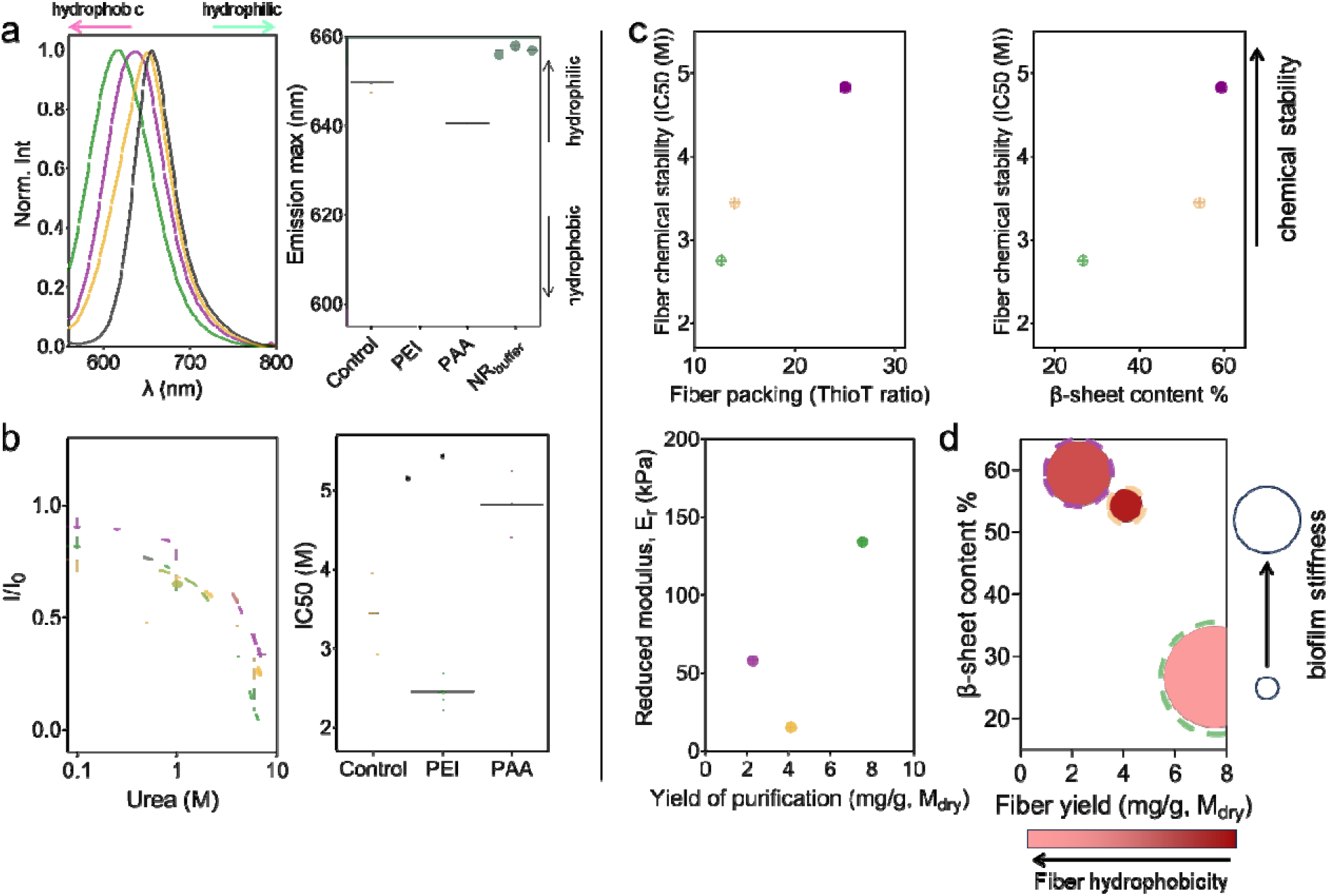
Hydrophobicity and chemical stability of the purified curli fibers. (a) Representative spectra of the Nile Red (NR) bound to the fibers emission (left panel). The NR in buffer spectrum was put as a reference of the probe in highest hydrophilic environment possible (grey line). Position of the emission peak of the fluorescence spectra of fibers stained with Nile Red (NR) (right panel). The shadowed areas in the extremes for hydrophobic and hydrophilic environments (pink and green, respectively). (b) Chemical stability curves of the purified fibers upon denaturation with increasing urea concentrations (0.1– 8⍰M) (left panel). The presence of the fibers was observed by ThioT fluorescence emission intensity. I0 corresponds to the ThioT emission when bound to fibers without urea in the solution. IC50 values for each curve in the left panel (right panel). IC50 corresponds to the urea concentration at which the ThioT intensity is 50% of the initial one. The statistical analysis was done with One-way ANOVA, post test used was Tukey’s test to compare each fiber against every fiber (p < 0.0001, **** | p < 0.001, *** | p < 0.01, ** |p < 0.05, * | ns = non-significant). (c) Relationship between fiber yield and properties such as structure and chemical stability (IC50) of the different curli fibers, and biofilm stiffness (reduced modulus, E_r_. The parameters taken as representatives for the fiber structure where ThoiT ratio (Figure 3b), and β-sheet content (Figure 3d, Table 1). The parameters for the biofilm stiffness were taken from Figure 2a. (d) Correlation matrix between fiber characteristics and biofilm stiffness. This plot describes the correlation between fiber yield with fiber hydrophobicty, fiber structure, and biofilm stiffness. The date represented in these plots was data taken from the experiments done in this study, in which mean values are represented, except for median values for the biofilm stiffness. For all experiments, N=3 independent purification batches of fibers grown in each condition.

The chemical stability of the fibers was studied upon incubation with increasing urea concentrations (Figure 4b). The loss of β-sheet structure of the fibers, monitored by ThioT emission, was considered as indicative of fiber denaturation.^7,8^ We calculated the urea concentration needed to reduce the ThioT emission by half (IC50); a decrease by half of ThioT emission can be interpreted as reducing the presence of curli fibers by half. Curli fibers from PAA-biofilms had the highest IC50 values reporting on the highest stability (5 ± 0.4 M), while fibers from biofilms grown on non-modified surfaces and PEI-grown biofilms had lower chemical stabilities (IC50 values of 3 ± 0.5, and 2 ± 0.2 M, respectively). Based on the results of the study of fiber structure by ThioT emission, and ATR-FTIR spectroscopy, there might be a correlation between the packing of the fibers and the β-sheet content with the stability of the fibers (Figure 4c). The higher the fiber packing (represented by the ThioT ratio in Figure 3b), and the β-sheet content, the more stable the fiber was against urea.

## Discussion

PEI and PAA have been studied as components for coatings in surfaces to avoid bacterial attachment, and biofilm formation of several bacteria types and strains.^22,23^ In this work, we deepen the knowledge on how agar surfaces coated with these polyelectrolytes influence *E. coli* W3110 biofilm’s macroscopic characteristics, and molecular properties. PEI-grown biofilms became 3 times smaller than biofilms grown on non-modified surfaces (Figure 1a-b, Figure 5). While their weight (M_dry_) was half the weight of control biofilms, their density was 1.15 times higher. In contrast, PAA-grown biofilms did not show significant differences when compared to biofilms grown on non-modified surfaces in the same experiments. PAA-grown biofilms were only 1.15 times smaller in area and in weight, while maintaining the same surface density as biofilms grown on non-modified surfaces (Figure 1a-b). A previous work from our group on another *E. coli* strain rendered similar results.^15^ Upon seeding, initial bacteria adhesion is affected by the substrate physicochemical characteristics mainly through interaction with their membrane.^24^ Moreover, the physicochemical characteristics of the agar substrate affect the spreading capacity of the biofilms, causing continuous or arrested spreading.^14^ These points could explain the differences observed in our data. Interestingly, PEI-grown biofilms yield almost twice as much curli as the biofilms grown on non-modified surfaces which correlated with a higher water uptake after biofilm dehydration (Figure 1e, Figure 5). A higher amount of curli amyloid fiber in the ECM might result in a more crosslinked extracellular mesh, which in turn would result in less irreversible changes of curli during drying as observed for *B. subtilis* biofilms.^25^ Further evidence of this would be the low fiber yield in PAA-grown biofilms, together with their low capacity of water uptake (Figure 1d-e).

**Figure 5.**
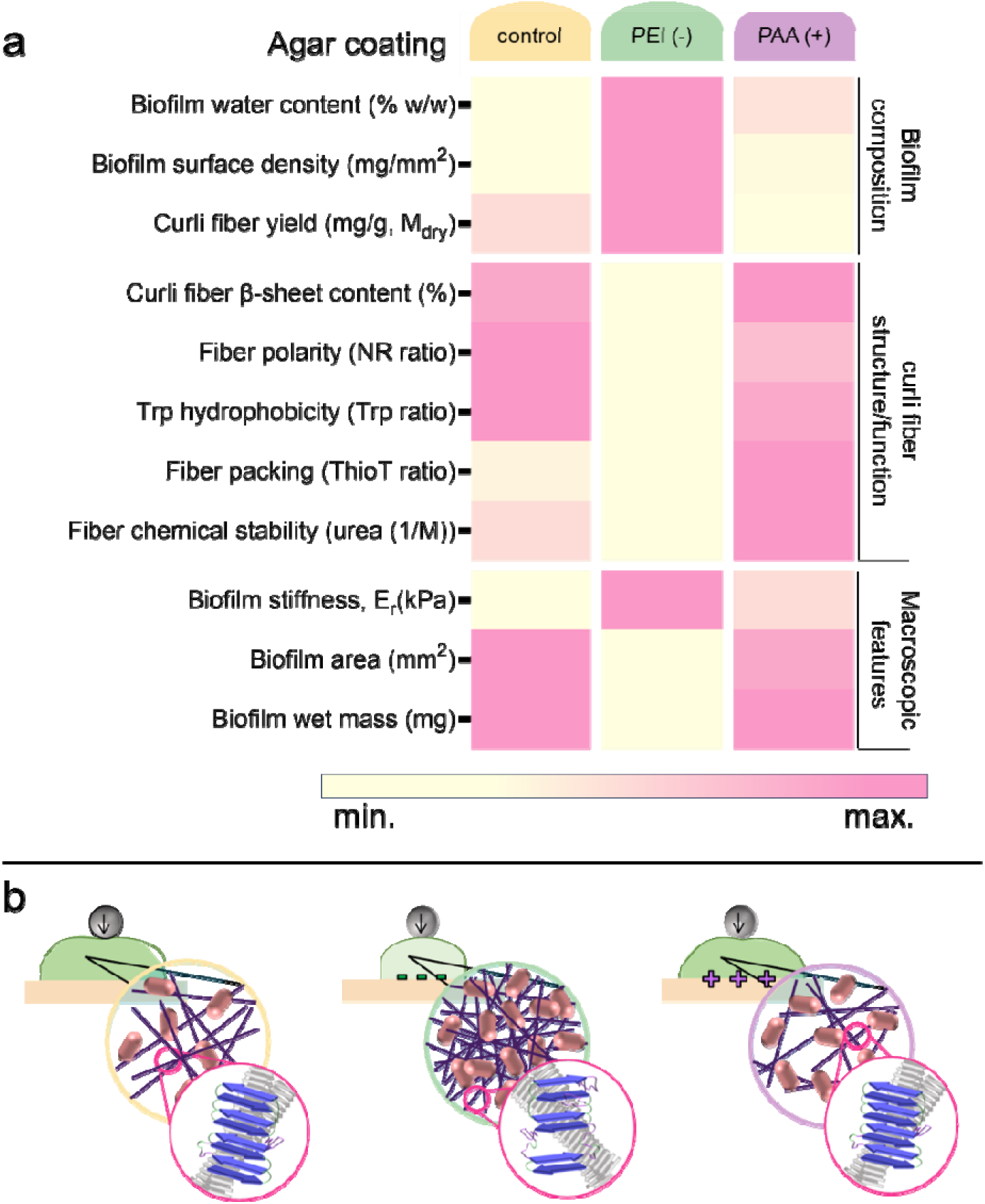
Summary of the relationship between the polyelectrolyte used to coat the salt-free LB agar and the macro- and microscopic features of the corresponding biofilms. (a) Table where the color shade represents qualitatively and comparatively the parameter value in the column of the left. (b) Sketch depicting the characteristics of biofilms grown on non-coated and coated surfaces.

A higher amount of curli amyloid fibers in the ECM could also explain the higher biofilms stiffness observed for PEI-grown biofilms despite their higher water content (Figure 2a-b, Figure 4c, and Figure 5). Amyloid fibers are known to be rigid robust proteinous structures, which largely contribute to biofilm stiffness.^10,20,26^ Their absence in the biofilm ECM decreases by 20-fold in mutants lacking these fibers.^10^ Nevertheless, PAA-grown biofilms which yielded half of the fibers biofilms grown on non-modified substrates were stiffer than the control biofilms. These results hint at the possibility that parameters other than fiber quantity might contribute to the biofilm emergent properties. For instance, PAA-grown biofilms produced fibers with a higher packing ratio, β-sheet content (Figure 3b, and Figure 3d) and chemical stability (Figure 4b, Figure 5), while PEI-grown biofilms produced fibers that are more hydrophobic (Figure 3c, and Figure 4a), but less structured as shown by their larger β-turn content over their β-sheet (Figure 3d). *In vitro* experiments done on a wide pH range of CsgA polymerization into curli fibers, showed no structure changes in the mature fibers.^27^ Yet, the structure differences observed in the curli amyloid fibers from biofilms grown on the different agar surfaces may result from how the CsgA monomers interact with the charged surfaces during fiber polymerization. Indeed, DeBenedictis et al. showed that the location and nature of the residues in the CsgA monomer sequence help discriminate the rearrangement of the flexible sections in the protein and what their interaction will be with charged surfaces.^28^

Previous experiments on biofilms showed that variations in water and nutrient content in the agar substrates resulted in biofilms with curli amyloid fibers of different structures.^4,19^ It has been reported that the fibrillation process of curli during biofilm formation would not always be homogenous especially in the interaction between monomers.^29^ Thus, changes observed in the structure of the curli fibers in this work might be related not only to the environmental changes due to the charged coatings, but also could be related to the changes that bacteria undergo in these environments. While this is something yet to be studied in details, previous experiments have touched upon this aspect.^4^ Fiber quantity and structure are important characteristics of amyloid fibers contributing to biofilm hydrophobicity,^10^ and mechanical properties (Figure 4d, Figure 5 and Figure S3).^4,10,19^ While bacteria exposed to positively-charged coatings aim to produce a higher amount of curli (forgoing their ordered structure), bacteria exposed to negatively-charged surfaces might aim to low amount of curli but enhance their biophysical properties (chemical stability and β-sheet content and packing) to produce a more robust matrix (Figure 4d). Nevertheless, a high amount of curli in the ECM could result in larger water uptake of the biofilm, fulfilling one of the main functions of the ECM (Figure 1e, and Figure S3). Interestingly, it seems that the biofilms grown on non-modified agar substrates rendered a balance of these features (Figure 4d, and Figure S3). These biofilms were able to render an ECM with high yield of fibers with enough structure (β-sheet content) and chemical stability, contributing to biofilms that, although not as stiff, had a higher water uptake per fiber yield ratio.

Until now, most of the studies on the charged surface-biofilm interaction have been focused on strategies to eradicate biofilms.^6,17^ Understanding the mechanisms of these interactions may open up new avenues into exploiting the surface-biofilm relationship for antibiofilm strategies, as well as farming biofilms as producers of bio-sourced materials (e.g. amyloid fibers). Biofilm ECM displays a complex organization, which attests for complex reactions to surfaces.^18,20,26^ In such a way, positively charged surfaces could serve to weaken and confine biofilm growth, but also to promote the production of curli amyloid fibers with specific biophysical properties, as tuned building blocks for specific biomaterials.^7,30,31^

## Conclusion

We explored how polyelectrolyte coatings applied on agar substrates influence *E. coli* biofilm macroscopic properties, but also the molecular structure of their amyloid-based ECM. The data indicated that the contribution of amyloid fibers to biofilm stiffness is given by a balance between quantity and quality (i.e. packing and β-sheet content). Further studies are needed to fully understand if the final structure of the amyloid fibers is given exclusively by their assembly conditions in the ECM environment during biofilm growth, or due to the metabolic state of the bacteria. Nevertheless, this work contributes to understanding the relationship between environmental conditions, structure and function of bacteria bio-sourced materials such as amyloid fibers. Such knowledge is precious for further developments of antimicrobial strategies as well as for the design and production of ELMs.^6,7,17^

## Experimental Section

### Agar preparation and polyelectrolyte coating

Salt-free agar plates (15⍰mm diameter) were prepared with 1.8% w/v of bacteriological grade agar−agar (Roth, 2266), supplemented with 1% w/v tryptone (Roth, 8952) and 0.5% w/v yeast extract (Roth, 2363). To prevent condensation in the plate after agar pouring, the plates were left to dry for 10 min with the lid open and 10 min with the lid partially open. Each agar plate was left to rest for 48 h before bacteria seeding. Polyelectrolytes solutions and coating were prepared following the work of Ryzhkov et al.^15^ 2 mg mL^−1^ solutions of branched polyethyleneimine (PEI; weak polycation, *M*_w_ = 25⍰000, Sigma-Aldrich), and of poly (acrylic acid) (PAA; weak polyanion, *M*_v_ = 450⍰000, Sigma-Aldrich) were prepared in 0.5 m NaCl. The solutions were sterilized using filtration and UV irradiation. Before bacteria seeding, each agar plate was divided in an imaginary 9 square-grid and a 50 μL drop of the polyelectrolyte solution was placed in each part of the grid to be coated, allowing for spontaneous spreading. The drops were left to dry for 40 minutes with the plate lid off. Parts of the grid dedicated to biofilm growth in control conditions were not coated.

### Bacterial strain and growth

The biofilm-forming bacterial strain *E. coli* K-12 W3110 was used throughout this study. A suspension of bacteria was prepared from a single colony and grown overnight in Luria–Bertani (LB) medium at 37 °C with shaking at 250 rpm. A 5 μL of bacterial suspension (OD600 ≈0.5 after 10× dilution) was seeded in each part of the grid (i.e. on the coated area), and the excess water was left to evaporate for 10 minutes with the plate lid off. Biofilms were grown for 5 days (≈120 h) inside an incubator at 28 °C and 30%RH.

### Biofilm imaging

Three biofilms per condition were imaged with a stereomicroscope (AxioZoomV.16, Zeiss, Germany) using the tiling function of the acquisition software (Zen 2.6 Blue edition, Zeiss, Germany). To estimate the biofilm size, 3 independent biofilms were measured at their equatorial line using the Fiji software.^13^ An average was then calculated for each growth condition.

### Biofilm water content, dry mass and water uptake

The water content and water uptake of the biofilms were determined by scraping 7 biofilms per condition from the respective agar substrates after 5 days of growth (∼120 h). Biofilms were placed in plastic weighing boats, and dried at 60°C for 3 h in an oven. Wet and dry masses (m_wet_, m_dry_) were determined before and after drying.^14^ To determine the water uptake (W_up_), we added Millipure water in excess (5 mL) to the biofilms harvested from each condition, covered them with aluminum foils to avoid evaporation and left overnight. The water excess was removed and the biofilm samples were weighed again (m_rewet_). The biofilms water content in each growth condition was estimated with **Eq. (1)**

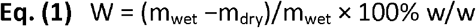

The percentage of water uptake of biofilms after rehydration (%W_up,w_) was determined with respect to biofilm initial wet mass as described in **Eq. (2)**

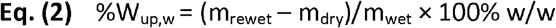

The water uptake per gram of dry biofilm (W_up,d_) was calculated with **Eq. (3)**

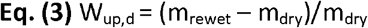

All procedures were carried out in four independent experiments.

### Micro-indentation on biofilms

*E. coli* biofilms of each strain were grown for 5 days on the same plate and used for the indentation experiments.^26^ A TI 950 Triboindenter (Hysitron Inc.) equipped with a conospherical tip (r=50 μm) was used to determine the load–displacement curves after calibration of the instrument in air. Ten measurements were performed in the central region of each biofilm. The distance between two measurement points was at least 250 μm in x and y directions and the depth of the indentation was between 10 and 25 μm, i.e. much less than biofilm the thickness (∼ 75 μm). Loading rates ranged from 20 μm/s, which translates to loading and unloading times of 10 s. The loading portion of all curves were fitted with a Hertzian contact model over and indentation range of 0 to 10 μm to obtain the reduced elastic modulus E_r_ .^4,19,26^

An apparent plasticity index (ψ’) was defined as **Eq. (5)**^20^

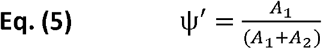

where A_1_ describes the area between the loading and unloading curves and A_2_ describes the area under the unloading curve. ψ’ spans between the values 0 – 1, where ψ’ = 1 characterizes a plastic behavior (irreversible deformation), and ψ’ = 0 indicates an elastic response (reversible deformation).

A subset of curves with a maximum indentation depth of 20 μm were selected for the analysis of the adhesion force (*F*_ad_)^26^ ; the minimum force measured during tip retraction.^20^

### Curli fiber purification and quantification

Fiber purification involved a similar process as reported in previous works.^3,4^ Briefly, a total of 27 biofilms (∼ 1g of biofilm material) were scraped from the surface of the substrates. Biofilms were blended five times on ice with an XENOX MHX 68500 homogenizer for 1min at 2-min intervals. The bacteria were pelleted by centrifuging two times at low speed (5000g at 4°C for 10min). A final concentration of NaCl 150 mM was added to the supernatant and the curli pelleted by centrifuging at 12.000g at 4°C for 10 minutes. The pellet was resuspended in 1mL of solution containing 10mM tris (pH 7.4) and 150mM NaCl, and incubated on ice for 30min before being centrifuged at 16.000g at 4°C for 10 minutes. This washing procedure was repeated thrice. The pellet was then resuspended in 1mL of 10mM tris solution (pH 7.4) and pelleted as described above (16.000g at 4°C for 10 minutes). The pellet was again suspended in 1mL of 10mM tris (pH 7.4) and centrifuged at 17.000g at 4°C for 10 minutes. This washing step was repeated twice. The pellet was then resuspended in 1mL of SDS 1% v/v solution and incubated for 30min. The fibers were pelleted by centrifuging at 19.000g at 4°C for 15min. The pellet was resuspended in 1mL of Milli-Q water. This washing procedure was repeated thrice. The last resuspension was done in 0.1mL of Milli-Q water supplemented with 0.02% sodium azide. The fiber suspension was stored at 4°C for later use. The protein concentration in monomeric units of the suspensions was determined by the absorbance from an aliquot incubated in 8M urea at 25°C for 2h, a treatment leading to complete dissociation of the fibrils as verified by Thioflavin T measurements.

### Transmission electron microscopy (TEM)

2μL drops of fiber suspension were adsorbed onto Formvar-coated carbon grids (200 mesh), washed with Milli-Q water, and stained with 1% (w/v) uranyl acetate. The samples were imaged in a JEOL-ARM F200 transmission electron microscope equipped with two correctors for imaging and probing. For the observations, we applied an acceleration voltage of 200 kV. The width of the purified fibers values in TEM images were measured using the scale tool of the GATAN GMS 3 software. To avoid subjectivity over 10 different images and over 10 identified fibers per field were used.

### Fluorescence spectroscopy: ThioT emission

Fiber samples were normalized to a 5μM CsgA monomer concentration (as previously described) for all fluorescence spectroscopy experiments. Corrected steady-state emission spectra were acquired with a FluoroMax®-4 spectrofluorometer (HORIBA). Spectra were recorded at 25°C using a 3-mm path cuvette (Hellma® Analytics). ThioT measurements were performed at final concentrations of 3 μM protein, 1 mM probe in Glycine buffer, pH 8.2, using λ_exc_ = 446 nm and spectral bandwidths of 10 nm.

### Attenuated total reflectance Fourier transform infrared spectroscopy (ATR-FTIR)

IR spectra were acquired on a spectrophotometer (Vertex 70v, Bruker Optik GmbH, Germany) equipped with a single reflection diamond reflectance accessory continuously purged with dry air to reduce water vapor distortions in the spectra. Fibers in Milli-Q water samples (∼10μL) were spread on a diamond crystal surface, dried under N_2_ flow to obtain the protein spectra. A total of 64 accumulations were recorded at 25°C using a nominal resolution of 4cm^−1^.

Spectra were processed using Kinetic software developed by Dr. Erik Goormaghtigh at the Structure and Function of Membrane Biology Laboratory, Université Libre de Bruxelles, Brussels, Belgium. After subtraction of water vapor and side chain contributions, the spectra were baseline corrected and area normalized between 1700 and 1600cm^−1^ (Figure S2). For a better visualization of the overlapping components arising from the distinct structural elements, the spectra were deconvoluted using Lorentzian deconvolution factor with a full width at the half maximum (FWHM) of 20 cm^−1^ and a Gaussian apodization factor with a FWHM of 30 cm^−1^ to achieve a line narrowing factor K = 1.5.^17^ Second derivative was performed on the Fourier self-deconvoluted spectra for band assignment. The bands identified by both procedures were used as initial parameters for a least square iterative curve fitting of the original IR band (K = 1) in the amide I’ region, using mixed Gaussian/Lorentzian bands. Peak positions of each identified individual component were constrained within ±2 cm^−1^ of the initial value.

### Fluorescence spectroscopy

Corrected steady-state emission spectra were acquired with a FluoroMax®-4 spectrofluorometer (HORIBA). Spectra were recorded at 25 °C using a 3-mm path cuvette (Hellma® Analytics). Thioflavin T(ThioT) measurements were performed at final concentrations of 3 μM protein, 1 mM probe in Glycine buffer, pH 8.2, using λ_exc_ = 446 nm and spectral bandwidths of 10 nm. Nile red (NR) measurements were performed at a final concentration of 7 μM protein, 7 mM probe in water, using λ_exc_ = 560 nm and spectral bandwidths of 10 nm. Intrinsic fluorescence spectra (5 μM protein) were acquired using λ_exc_= 280 nm and 5/5 nm slit bandwidths. 5 μM solution of soluble Trp (N-Acetyl-L-tryptophan, NATA) was included as a reference of the emission spectrum of a fully exposed Trp.

Fiber polarity is defined as the ratio between the position of the emission maximum of the nile red (NR) in buffer and the position of the emission maximum of the NR bound to each fiber

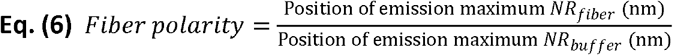

A polarity value of 1.00 indicates highly polar fibers.

Trp hydrophobicity is defined as the ratio between the position of the emission maximum of the Trp population in the fibers and the position of the emission maximum of soluble Trp (NATA) in buffer (as reference for the position of the emission maximum when we have the highest exposure to the solvent).

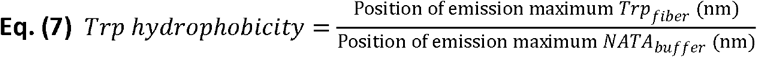

A hydrophobicity value of 1.00 indicates more exposure of the Trp to the solvent.

### Chemical stability assay

To test the chemical stability of the protein fibers, 5 μM samples were prepared by incubation of increasing urea concentration (0 – 8 M) and left for 2 h at room temperature to ensure equilibrium. ThioT was then added in a final concentration of 1 mM and fluorescence emission spectra of the samples were acquired under excitation at λ_exc_ = 446 nm and spectral bandwidth of 5 nm. Emission of the ThioT in urea was done and no significant signal was detected.

The IC50 value was estimated by fitting the data to a linear regression (Y = a * X + b) and then calculating IC50 = (0.5 - b)/a.

### Statistical analysis

For each experiment, 3 to 4 fiber solutions were used, where each solution came from different fiber purification batches. For each purification, 27 biofilms were cultured in each of the 4 growth conditions tested (i.e. on salt-free LB-agar plate containing 0.5%, 1.0%, 1.8% and 2.5% agar respectively). For each batch of biofilm culture, the different samples of fibers obtained for the 4 conditions were treated simultaneously (or in consecutive days) to avoid variability due to unavoidable slight variations in the implementation of the protocols (e.g. temperature and humidity in the laboratory during agar preparation and/or biofilm seeding).

For statistical analysis, a Shapiro Wilk test was used to check for data normality. For data with no normal distribution, Kruskal-Wallis non-parametric test was performed. For data with normal distribution, a One-way ANOVA test was carried out. Mechanical properties data was analyzed using a Mann-Whitney U test. Unless otherwise stated in the caption, Dunn’s post-test for multiple comparisons were done with respect to the 1.8 % salt-free LB-agar condition, considered as the standard seeding condition. Details of each test are described in the legend of the figures.

## Supporting Information

The following files are available free of charge:

- Biofilm morphology: Figure S1
- Biofilm mechanical properties: Figure S2
- Fiber and biofilm characteristics correlation matrix: Figure S3

## Data Availability Statement

The data that support the findings of this study are available from the corresponding authors upon reasonable request.

## AUTHOR INFORMATION

### Author Contributions

The manuscript was written through contributions of all authors. All authors have given approval to the final version of the manuscript. The authors declare no competing interests.

## ACKNOWLEDGMENTS

M.S. acknowledges support from the Max Planck Queensland Centre on the Materials Science for Extracellular Matrices. The authors also thank Christine Pilz-Allen for her technical support in the laboratories, Peter Werner and Heike Runge for their help in doing the transmission electronic microscopy experiments. The authors are also grateful to Regine Hengge (HU Berlin) for providing the *E. coli* strain W3110 and to Eric Goormaghtigh from the SFMB group at the Université Libre de Bruxelles for providing the Kinetics Software. Open Access funding enabled and organized by Projekt DEAL (Max Planck Society).

## Notes

### Competing Interest Statement

The authors have declared no competing interest.

