## Supplementary Information for "Charged agar surfaces affect *E. coli* biofilm properties by balancing curli amyloid quantity and quality"

#### Supplementary Information Table of Contents

| Biofilm morphology | Figure S1 |
| --- | --- |
| Biofilm mechanical properties | Figure S2 |
| Fiber and biofilm characteristics correlation matrix | Figure S3 |

### 1- Biofilm morphology

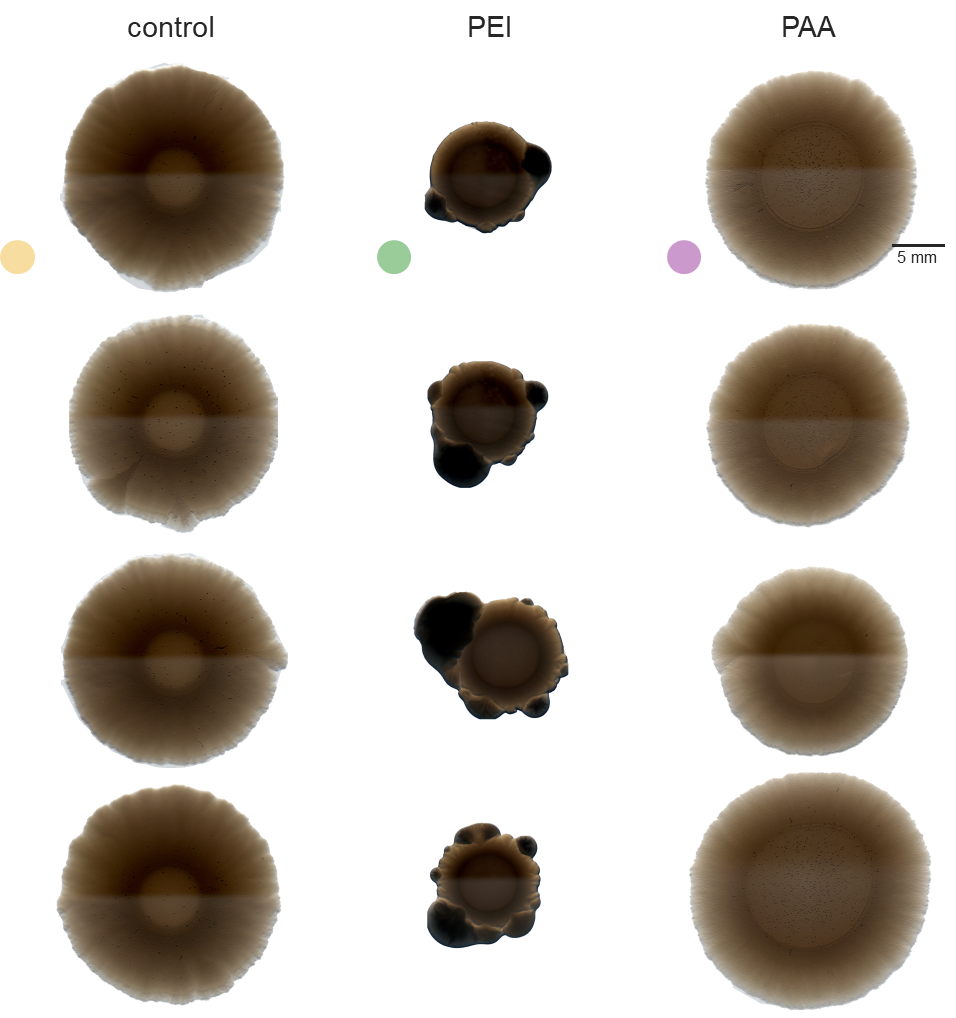

Figure S 1 Biofilm morphology. Stereomicroscopies of the variability of the morphology of *E. coli* biofilms grown under the different conditions studied. Scale bar = 5 mm

### 2- Biofilm mechanical properties

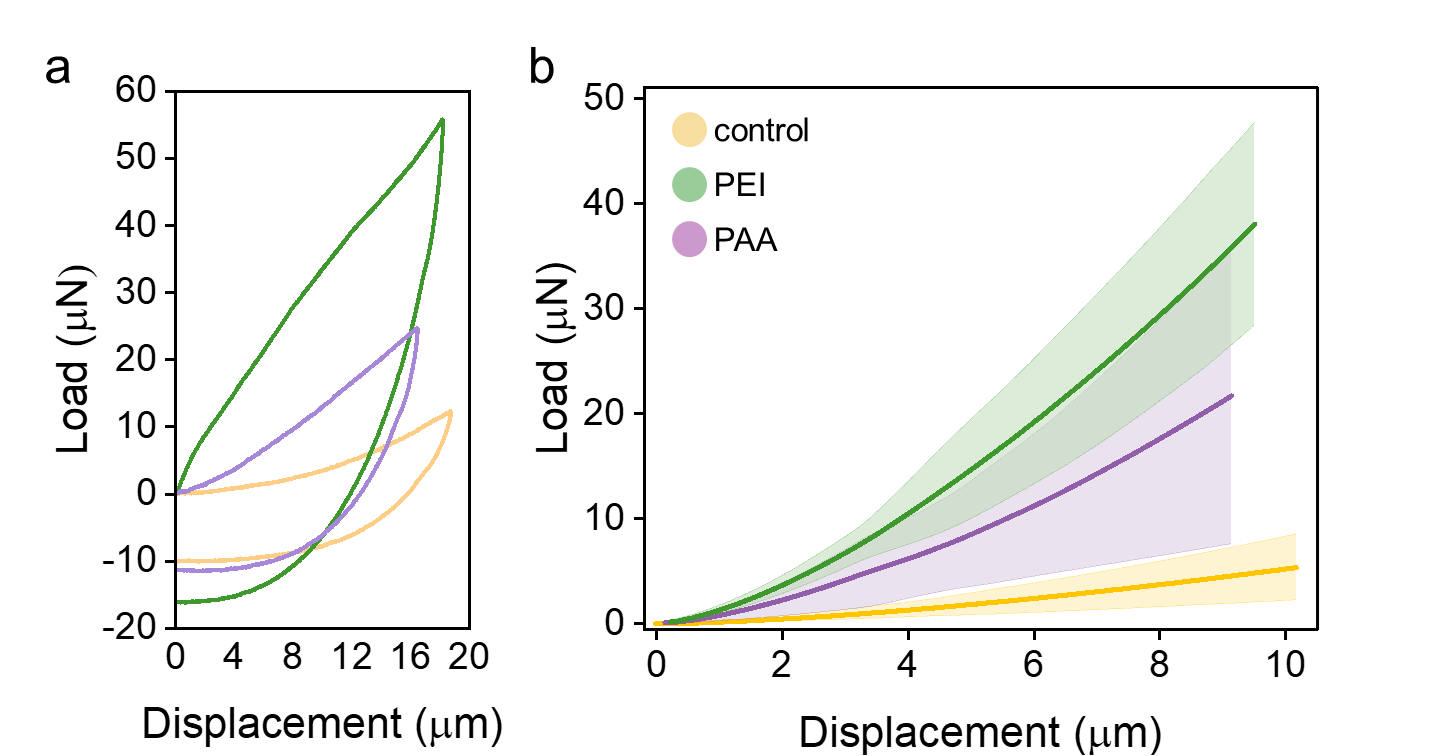

Figure S 2 Representative loading curves of microindentation experiments. (a) Representative load displacement curves when indenting the biofilm surface with a spherical tip indentation with a diameter of 50 μm. (b) Zoom in on part of the loading curves used for fitting a Hertzian contact model. Fitting curve ranges for reduced modulus estimation of all samples studied. Considering the size of the tip (conospherical tip, r=50 µm) and thickness of the biofilms (̴100 µm), the displacement range taken into account for the reduced modulus was ~10 μm.

### 3- Fiber and biofilm characteristics correlation matrix

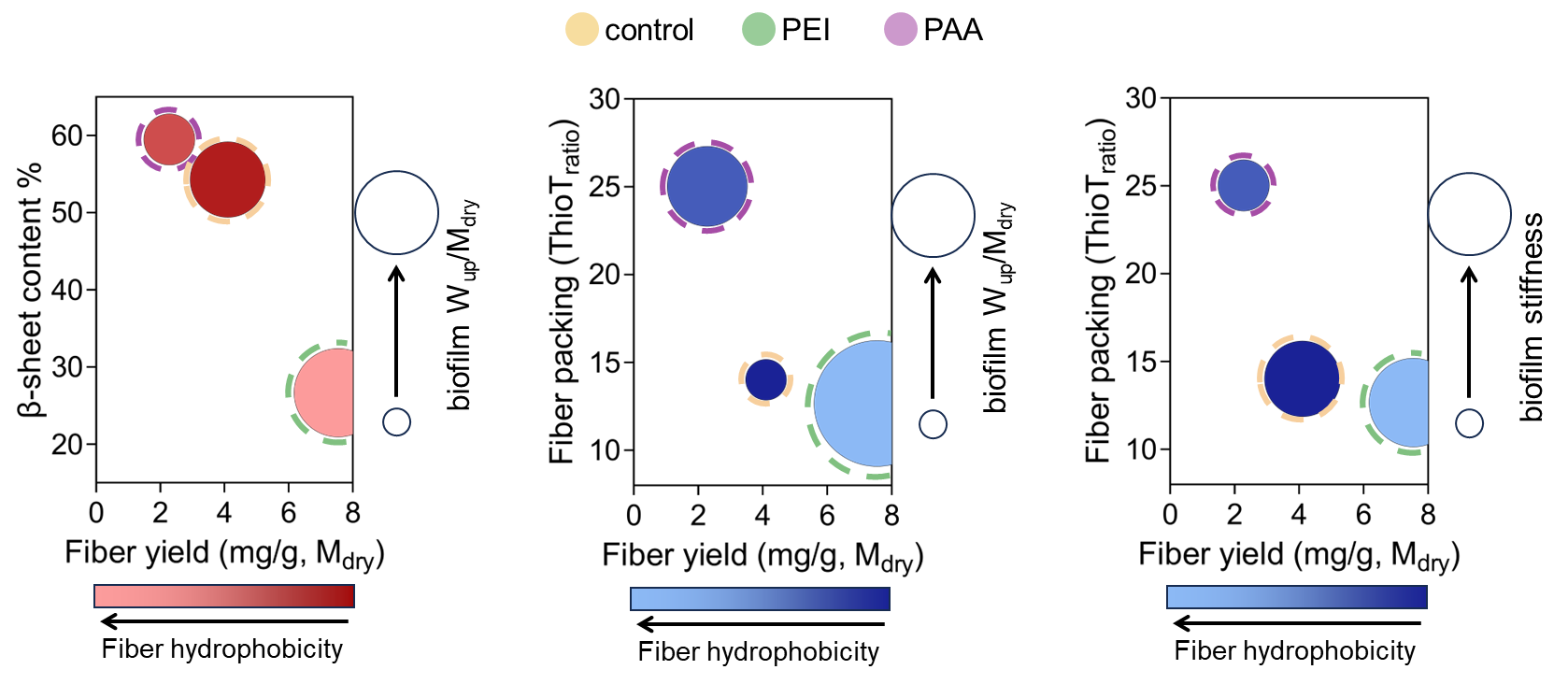

Figure S 3 Correlation matrix between fiber (fiber β-sheet content or fiber packing, fiber yield per M_dry_, fiber and fiber hydrophobicity) and biofilm characteristics (biofilm stiffness or biofilm water uptake). The data represented in these plots was data taken from the experiments done in this study, in which mean values are represented, except for median values for the biofilm stiffness. Red plots depict fiber β-sheet content in the Y-axis, while blue plots depict fiber packing (ThioT_ratio_) in the Y-axis

1. *Current address: Departamento de Química, Catedra de Química Biológica, Facultad de Ciencias Exactas, Físicas y Naturales, Universidad Nacional de Córdoba, Córdoba 5000, Argentina.*

   *Consejo Nacional de Investigaciones Científicas y Técnicas (CONICET), Instituto de Investigaciones Biológicas y Tecnológicas (IIBYT), Córdoba 5000, Argentina.*
